# MRI-BASED DEEP LEARNING METHOD FOR DETERMINING METHYLATION STATUS OF THE O_6_-METHYLGUANINE–DNA METHYLTRANSFERASE PROMOTER OUTPERFORMS TISSUE BASED METHODS IN BRAIN GLIOMAS

**DOI:** 10.1101/2020.05.30.124230

**Authors:** Chandan Ganesh Bangalore Yogananda, Bhavya R. Shah, Sahil S. Nalawade, Gowtham K. Murugesan, Frank F. Yu, Marco C. Pinho, Benjamin C. Wagner, Bruce Mickey, Toral R. Patel, Baowei Fei, Ananth J. Madhuranthakam, Joseph A. Maldjian

**Affiliations:** Advanced Neuroscience Imaging Research (ANSIR) Lab, Department of Radiology, University of Texas Southwestern Medical Center, Dallas, Texas; Department of Neurological Surgery, University of Texas Southwestern Medical Center, Dallas, Texas; Department of Bioengineering, University of Texas at Dallas, Richardson, Texas

## Abstract

**PURPOSE:** Methylation of the O_6_-Methylguanine-DNA Methyltransferase (MGMT) promoter results in epigenetic silencing of the MGMT enzyme and confers an improved prognosis and treatment response in gliomas. The purpose of this study was to develop a deep-learning network for determining the methylation status of the MGMT Promoter in gliomas using T2-w magnetic resonance images only.

**METHODS:** Brain MRI and corresponding genomic information were obtained for 247 subjects from The Cancer Imaging Archive (TCIA) and The Cancer Genome Atlas (TCGA). 163 subjects had a methylated MGMT promoter. A T2-w image only network (MGMT-net) was developed to determine MGMT promoter methylation status and simultaneous single label tumor segmentation. The network was trained using 3D-Dense-UNets. Three-fold cross-validation was performed to generalize the networks’ performance. Dice-scores were computed to determine tumor segmentation accuracy.

**RESULTS:** MGMT-net demonstrated a mean cross validation accuracy of 94.73% across the 3 folds (95.12%, 93.98%, and 95.12%, standard dev=0.66) in predicting MGMT methylation status with a sensitivity and specificity of 96.31% ±0.04 and 91.66% ±2.06, respectively and a mean AUC of 0.93 ±0.01. The whole tumor segmentation mean Dice-score was 0.82 ± 0.008.

**CONCLUSION:** We demonstrate high classification accuracy in predicting the methylation status of the MGMT promoter using only T2-w MR images that surpasses the sensitivity, specificity, and accuracy of invasive histological methods such as pyrosequencing, methylation-specific PCR, and immunofluorescence methods. This represents an important milestone toward using MRI to predict glioma histology, prognosis, and response to treatment.

## INTRODUCTION

O_6_-methylguanine-DNA methyltransferase (MGMT) promoter methylation is a molecular biomarker of gliomas that has prognostic and therapeutic implications. Unlike isocitrate dehydrogenase (IDH) mutations and 1p/19q co-deletions, MGMT promoter methylation is an *epigenetic* event. Epigenetic events are functionally relevant but do not involve a change in the nucleotide sequence. Therefore, while MGMT promoter methylation is an important prognostic marker, it does not define a distinct subset of gliomas. MGMT is a DNA repair enzyme that protects normal and glioma cells from alkylating chemotherapeutic agents. The methylation of the MGMT promoter is an example of epigenetic *silencing* which results in loss of function of the MGMT enzyme and its protective effect on glioma cells. The survival benefit incurred by MGMT promoter methylation in patients treated with temozolomide (TMZ) was determined in 2005.^1^ Subsequent work by Stupp et al. has shown that in patients who received both radiation and temozolomide, *MGMT* promoter methylation improved median survival as compared to patients with unmethylated gliomas (21.7 months vs 12.7 months).^2,3^ Long-term follow-up from that initial study has further substantiated the survival benefit.^2,3^ As a result, determining MGMT promoter methylation status is an important step in predicting survival and determining treatment.

Currently, the only reliable way to determine MGMT promoter methylation status requires analysis of glioma tissue obtained either via an invasive brain biopsy or following open surgical resection. Surgical procedures carry the risk of neurologic injury and complications. Therefore, considerable attention has been dedicated to developing non-invasive, image-based diagnostic methods to determine MGMT promoter methylation status. A meta-analysis of MRI features demonstrated that glioblastomas with methylated MGMT promoters were associated with less edema, high apparent diffusion coefficient (ADC), and low perfusion. However, the summary sensitivity and specificity of these clinical features was only 79% and 78% respectively.^4^ Recent advances in deep-learning methods have not only made non-invasive, image-based molecular profiling possible but also highly accurate. Our group has previously demonstrated highly-accurate, MRI-based, voxel-wise deep-learning networks for determining IDH-classification and 1p/19q co-deletion status using only T2-w MR images.^5,6^ The benefits of using T2-w images are that they are routinely acquired, they can be obtained quickly, and high quality T2-w images can even be obtained in the setting of motion degradation. Because MGMT promoter methylation in gliomas is such an important biomarker, the purpose of this study was to develop a highly accurate, fully automated deep-learning 3D network for MGMT promoter methylation status determination using T2-w images only.

## MATERIAL & METHODS

### Data and Pre-processing

Multi-parametric MR Images of brain glioma patients were obtained from the TCIA (The Cancer Imaging Archive) database.^7,8^ The genomic information was obtained from both TCGA (the cancer genome atlas) and TCIA databases.^7,9,10^ Subject datasets were screened for the availability of pre-operative MR images, T2-w images and known MGMT promoter status. The final dataset of 247 subjects included 163 methylated cases and 84 unmethylated cases. TCGA subject IDs, MGMT status, and tumor grade are listed in Table 1 of the supplementary data.

**Table 1:** Cross-validation results.

| **Fold Description** | **MGMT-net** |  |  |
| --- | --- | --- | --- |
| **Fold Number** | **% Accuracy** | **AUC** | **Dice-score** |
| Fold 1 | 95.12 | 0.9574 | 0.8140 |
| Fold 2 | 93.98 | 0.8978 | 0.8165 |
| Fold 3 | 95.12 | 0.9390 | 0.8291 |
| **AVERAGE** | **94.73 +/- 0.66** | **0.93 +/- 0.03** | **0.82 +/- 0.008** |

Tumor masks for 179 subjects were available through previous expert segmentation.^5,11,12^ Tumor masks for the remaining 68 subjects were generated by our previously trained 3D-IDH network.^5^ These tumor masks were used as ground truth for tumor segmentation in the training step. Ground truth whole tumor masks for methylated and unmethylated MGMT promoter type were labelled with 1s & 2s respectively (Figure 1). Data preprocessing steps included (a) ANTs affine co-registration^13^ to the SR124 T2 template^14^, (b) skull stripping using the Brain Extraction Tool (BET)^15^ from FSL^15–18^, (c) Removing RF inhomogeneity using N4BiasCorrection^19^, and (d) normalizing intensity to zero-mean and unit variance. The pre-processing took less than 5 minutes per dataset.

**Figure 1:**
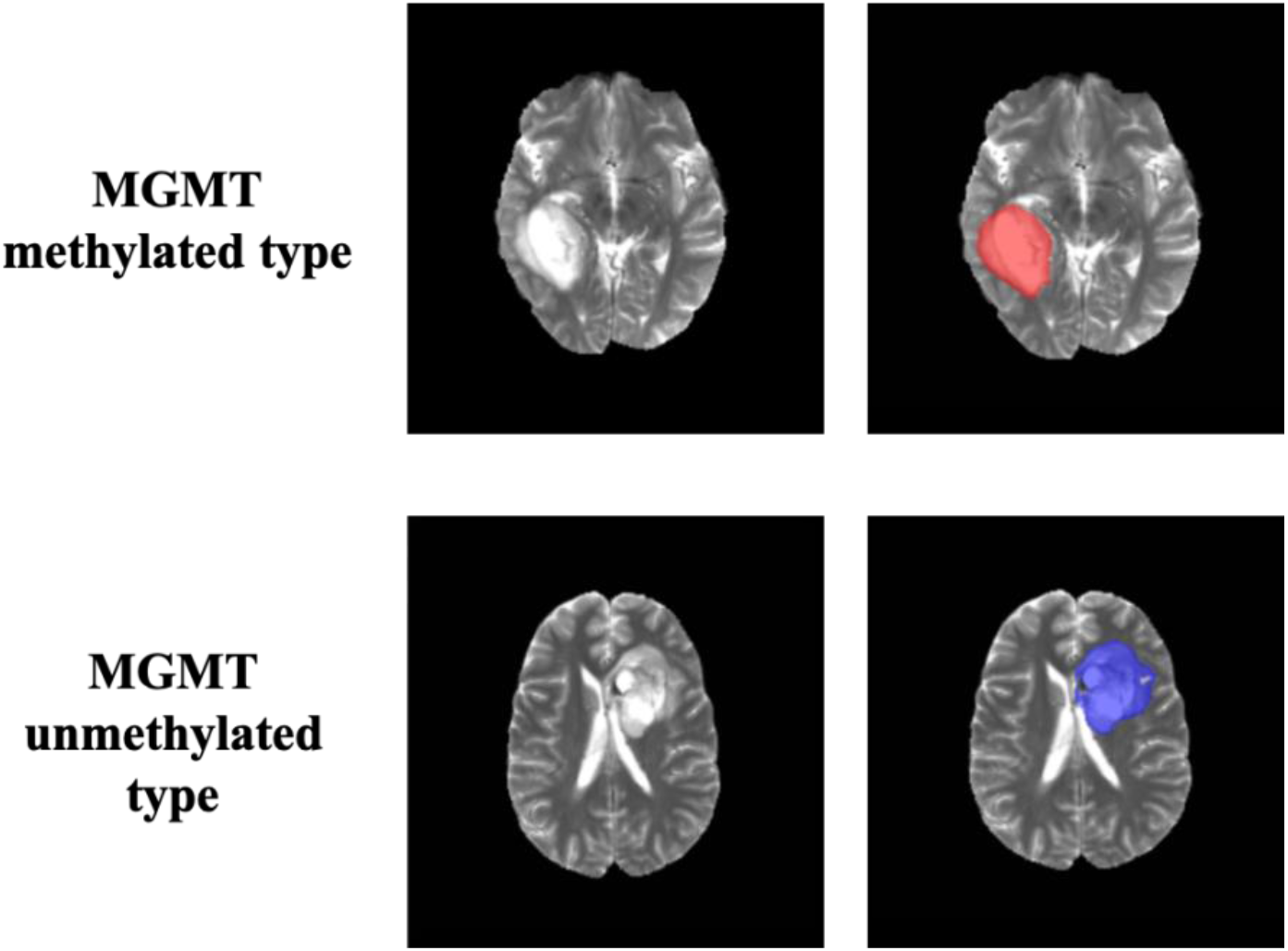
Ground truth whole tumor masks. Red voxels represent methylated MGMT promoter status (values of 1) and blue voxels represent unmethylated MGMT promoter status (values of 2). The ground truth labels have the same MGMT promoter status for all voxels in each tumor.

### Network Details

Transfer learning for MGMT promoter status determination was implemented using our previously trained 3D-IDH network.^5^ The decoder part of the network was fine-tuned for a voxel-wise dual-class segmentation of the whole tumor with Classes 1 and 2, representing methylated & unmethylated MGMT promoter type, respectively. The network architecture is shown in Figure 2B. A detailed schematic of the network is provided in the supplemental material section.

**Figure 2:**
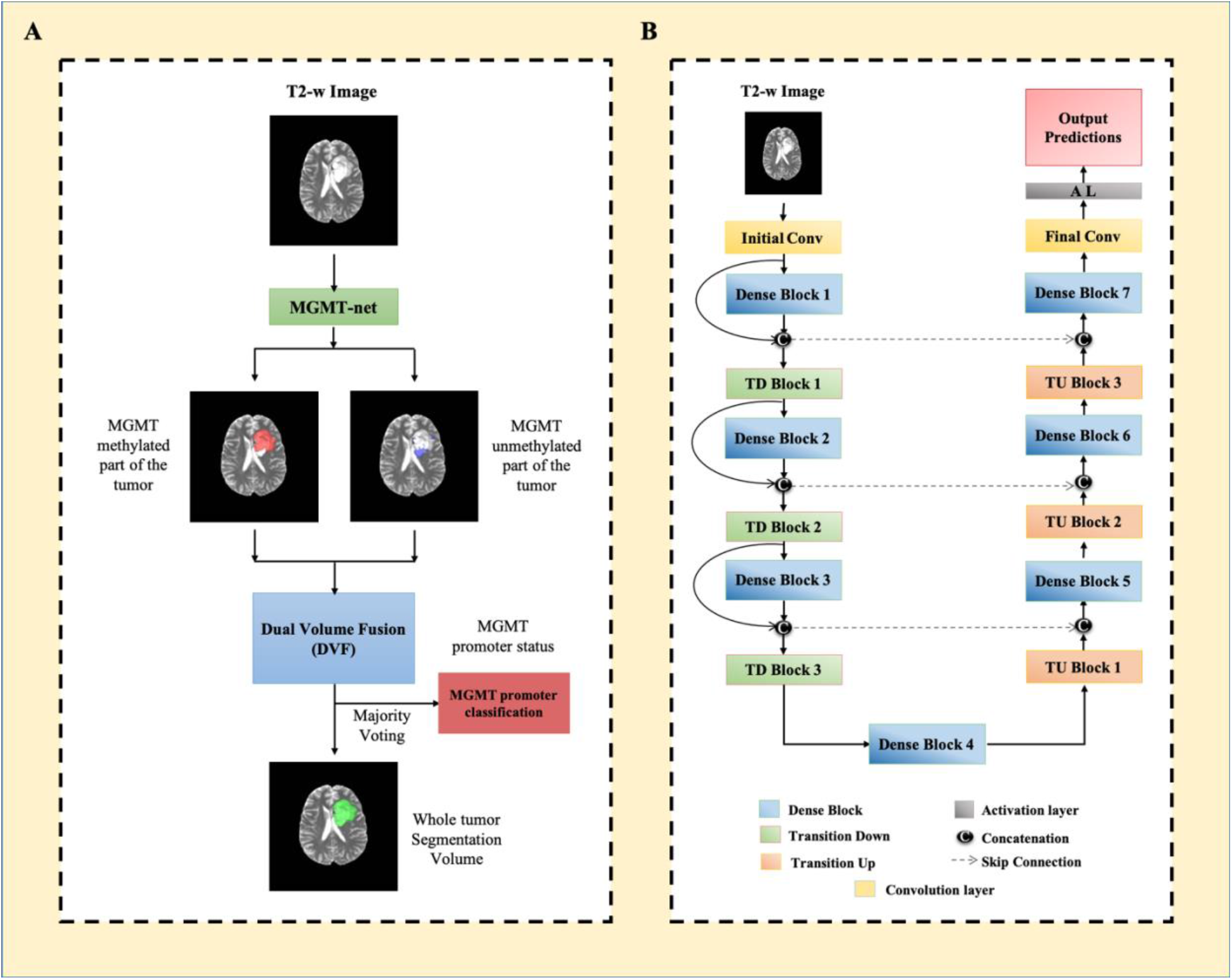
(A) MGMT-net overview. Voxel-wise classification of MGMT promoter status is performed to create 2 volumes (methylated and unmethylated MGMT promoter). Volumes are combined using dual volume fusion to eliminate false positives and generate a tumor segmentation volume. Majority voting across voxels is used to determine the overall MGMT promoter status. (B) Network architecture for MGMT-net. 3D-Dense-UNets were employed with 7 dense blocks, 3 transition down blocks, and 3 transition up blocks.

### Network Implementation and Cross-validation

To generalize the network’s performance, a 3-fold cross-validation was performed. The dataset of 247 subjects was randomly shuffled and distributed it into 3 groups (approximately 82 subjects for each group). Group 1 had 82 subjects (54 methylated, 28 unmethylated), Group 2 had 83 subjects (55 methylated, 28 unmethylated), and Group 3 had 82 subjects (54 methylated, 28 unmethylated). The 3 groups were alternated between training, in-training validation and held-out testing groups such that, each fold of the cross-validation was a new training phase based on a unique combination of the 3 groups. The network uses the in-training validation dataset to evaluate its learning after each training round and update model parameters to improve performance. However, the network performance is only reported on the held-out testing group for each fold as it is never seen by the network. The group membership for each cross-validation fold is listed in Table 1 of the supplementary data.

Seventy-five percent overlapping patches were extracted from the training and in-training validation subjects. To prevent the problem of data leakage, no patch from the same subject was mixed with the training, in-training validation or testing datasets.^20,21^ Data augmentation steps included horizontal & vertical flipping, random & translational rotation, addition of salt & pepper noise, addition of Gaussian noise, and projective transformation. Additional data augmentation steps included, down-sampling images by 50% and 25% (reducing the voxel resolution to 2mm^3^ and 4mm^3^). This provided a total of approximately 300,000 patches for training and 300,000 patches for in-training validation for each fold. The networks were implemented using Tensorflow^22^ backend engine, Keras^23^ python package, and an Adaptive Moment Estimation optimizer (Adam).^24^ The initial learning rate was set to 10^−5^ with a batch size of 15 and maximal iterations of 100.

MGMT-net outputs two segmentation volumes (V1 and V2), which are combined to generate the voxel-wise prediction of methylated & unmethylated MGMT promoter tumor voxels, respectively. The two volumes are fused and the largest connected component (3D-connected component algorithm in MATLAB^(R)^) is obtained as the single tumor segmentation map. Majority voting over the voxel-wise classes of methylated or unmethylated type provided a single MGMT promoter classification for each subject. Tesla V100s, P100, P40 and K80 NVIDIA-GPUs were used to implement the networks. This MGMT promoter determination process is fully automated, and a tumor segmentation map is a natural output of the voxel-wise classification approach.

### Statistical Analysis

Statistical analysis of the network’s performance was performed in MATLAB^(R)^ and R. Network accuracies were evaluated using majority voting (*i.e.* voxel-wise cutoff of 50%). The accuracy, sensitivity, specificity, positive predictive value (PPV), and negative predictive value (NPV) of the model for each fold of the cross-validation procedure were calculated using this threshold. Receiver Operating Characteristic (ROC) curves for each fold were generated separately. Dice-scores were calculated to evaluate the tumor segmentation performance of the networks. The Dice-score calculates the spatial overlap between the ground truth segmentation and the network segmentation.

## RESULTS

The network achieved a mean cross-validation testing accuracy of 94.73% across the 3 folds (95.12%, 93.98%, and 95.12%, standard dev=0.66). Mean cross-validation sensitivity, specificity, PPV, NPV and AUC for MGMT-net was 96.31% ±0.04, 91.66% ±2.06, 95.74% ±0.95, 92.76% ±0.15 and 0.93 ±0.03 respectively. The mean cross-validation Dice-score for tumor segmentation was 0.82 ± 0.008. The network misclassified 4 cases for fold one, 5 cases for fold two, and 4 cases for fold three (13 total out of 247 subjects). Six subjects were misclassified as unmethylated, and 7 as methylated.

### ROC analysis

The ROC curves for each cross-validation fold for the network is provided in Figure 3. The network demonstrated very good performance with high sensitivities and specificities

**Figure 3:**
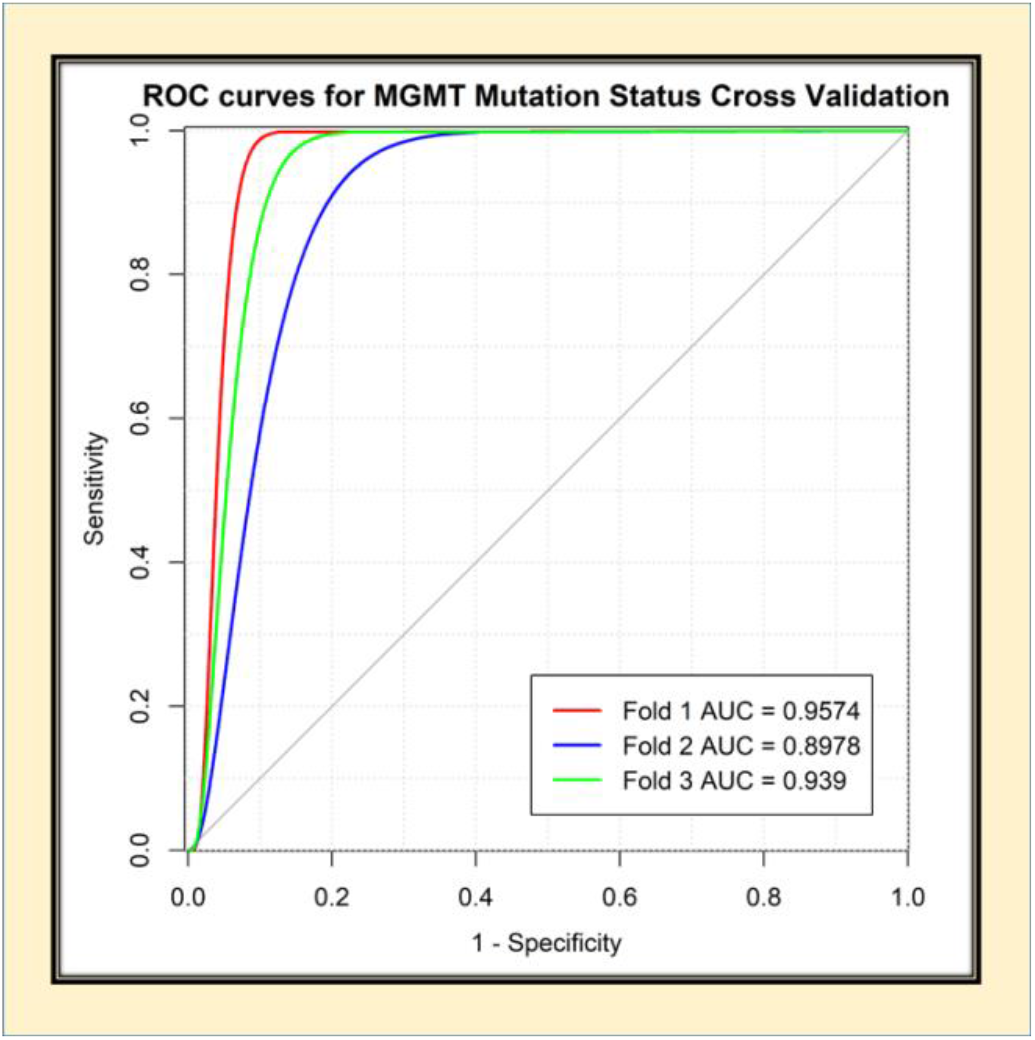
ROC analysis for MGMT-net. Separate curves are plotted for each cross-validation fold along with corresponding AUC value.

### Voxel-wise classification

The network is a voxel-wise classifier with the tumor segmentation map being a natural output. Figures 4A and 4B show examples of the voxel-wise classification for a methylated, and unmethylated MGMT promoter type respectively. The volume fusion procedure was effective in removing false positives and improving the dice-scores by approximately 6%. We also computed the voxel-wise accuracy for the network. The mean voxel-wise accuracies were 81.68% ±0.02 for methylated type and 70.83% ±0.04 for unmethylated type.

**Figure 4:**
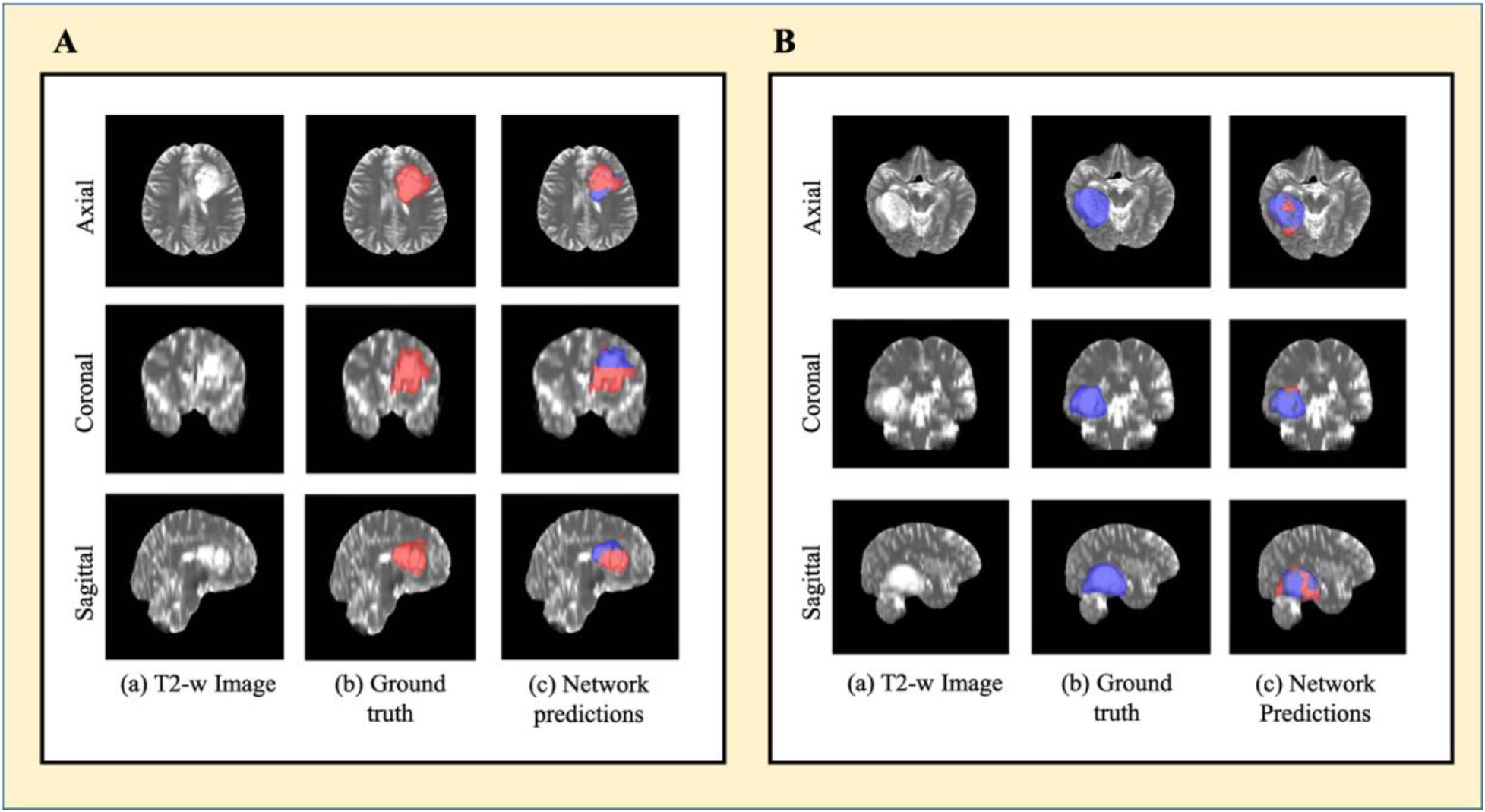
(A) Example of voxel-wise segmentation for a tumor with a methylated MGMT promoter. Native T2-w image (a). Ground truth segmentation (b). Network output after DVF (dual volume fusion) (c). Red voxels correspond to MGMT methylated class and blue voxels correspond to MGMT unmethylated class. (B) Example of voxel-wise segmentation for a tumor with an unmethylated MGMT promoter. The sharp borders visible between methylated and unmethylated type result from the patch-wise classification approach.

### Training and segmentation times

Fine-tuning the network took approximately one week. The trained network took approximately three minutes to segment the whole tumor and determine the MGMT status for each subject.

## DISCUSSION

We developed a fully-automated, highly accurate, deep-learning network for determining the methylation status of the MGMT promoter that outperforms previously reported algorithms.^25–27^ Our network is able to determine MGMT promoter methylation status from T2-w images alone. This obviates failure from image acquisition artifacts and makes clinical translation straightforward because T2-w images are routinely obtained as part of standard clinical brain MRI. Previous approaches have required multi-contrast input which can be compromised due to patient motion from lengthier examination times, and the need for gadolinium contrast. Obviating the need of intravenous contrast makes our algorithm applicable to patients with contrast allergies and renal failure. When compared to previously published algorithms, our methodology is fully automated and utilizes minimal preprocessing. The time required for MGMT-net to segment the whole tumor and predict the MGMT promoter methylation status for one subject is approximately 3 minutes on a K80 or P40 NVIDA-GPU.

The higher performance achieved by our network compared to previous image-based classification studies can be explained by several factors. The dense connections in our 3D network architecture are easier to train, carry information from the previous layers to the following layers, and can reduce over-fitting.^28, 29^ 3D networks also interpolate between slices to maintain inter-slice information more accurately. The Dual Volume Fusion (DVF) post-processing step improved the dice-scores by approximately 6% by eliminating extraneous voxels not connected to the tumor. Our approach also uses voxel-wise classifiers and provides a classification for each voxel in the image. This provides a simultaneous single-label tumor segmentation. The cross validation single label whole tumor segmentation performance for the MGMT network provided excellent Dice-scores of 0.82 +/− 0.008.

The ability to determine MGMT promoter methylation status based on MR images alone is clinically significant because it helps determine whether the glioma will be susceptible to temozolomide (TMZ). Alkylating agents such as TMZ, damage DNA by methylating the oxygen at position 6 of the guanine nucleotide (O_6_-methylguanine). The process by which many DNA repair enzymes remove O_6_-methylguanine, results in DNA breaks culminating in cell death. However, MGMT works differently by restoring the normal guanine residue and rescuing the glioma cell. Therefore, MGMT activity leads to resistance to therapy. Methylation of the MGMT promoter leads to inactivation of MGMT and loss of resistance of glioma cells to alkylating agents. The MGMT protein is encoded on the long arm of chromosome 10 at position 26 (10q26). Transcription of the MGMT gene is regulated by several promoters.^29^

Although incompletely understood, at least nine specific regions within the promoter’s gene determine whether a cell will express or not express MGMT.^29^ However, some regions have been shown to be more important for loss of MGMT expression.^30^ In the clinical setting, methods for determining MGMT methylation focus on these regions in the promoter’s gene. The four most prevalent methods to detect MGMT methylation are: immunohistochemistry (IHC), pyrosequencing (PYR), quantitative methylation-specific PCR (qMSP), and methylation-specific PCR (MSP). PYR is considered the theoretical “gold-standard” but is not readily available, and although it is quantitative, there is no agreement on what “cut-off” values to use when determining MGMT promoter methylation status.^30^ Therefore, although it is not quantitative, MSP is the most widely used method.^31^ Additionally, most centers perform MGMT methylation detection on formalin-fixed or paraffin embedded tissue specimens. These methods have several limitations. Evaluating multiple different methylation sites is technically challenging on a single tissue specimen.^31^ Tumor heterogeneity poses a substantial limitation for these methods because sampling-bias can lead to inaccurate determinations. The presence of hemorrhage, necrosis, or nonmalignant cells contaminate the specimen^31^. Therefore, some institutions mandate that at least 50% of the sample to be analyzed contains tumor cells. Prior to PCR, several tissue processing steps are required. Bisulfite treatment is the most critical step because it will produce the modified DNA that will be used for PCR; however, it also degrades the amount of DNA available and incomplete treatments can lead to false-positive results.^31^ The reported sensitivity and specificity of MSP is 91% and 75% respectively, while the reported sensitivity and specificity of PYR is 78% and 90%.^32^

Our non-invasive, MRI based deep learning algorithm outperformed these methods with a sensitivity and specificity of 96.3% and 91.6% respectively. The overall determination of MGMT promoter methylation status is based on the majority of voxels in the tumor. Given the variability in the cut-off values for pyrosequencing-based detection, we performed a Youden’s statistical index analysis to determine if the optimal cut-off for our deep learning algorithm was different than majority voting (>50%). The analysis demonstrated that maximum accuracy, sensitivity, specificity, PPV, and NPV were obtained at an optimal cut-off of 50%, the same as majority voting.

Our algorithm was trained on ground-truth obtained from the TCGA database. The TCGA uses Infinium assays to determine MGMT promoter methylation status.^32–34^ Infinium assays are an immunofluorescence method that uses next generation high-throughput microchip arrays and probes. While these methods have been reported to be more sensitive and specific than the most widely available clinical assays, these methods require pre-existing probes to detect specific methylation sites.^34^ The sensitivity and specificity values change depending on the probe and analytical model used to interpret the results.^34^ The sensitivities for the best probes range from 87.5-90.6% while the specificity is 94.4%.^34^ The overall accuracy of these probes with an optimized analytical model ranges from 91.24%-93.6%.^34^ Tissue based methods for determining MGMT promoter methylation status remain a complex, multi-step process that is susceptible to failure and inaccuracy even after an adequate tissue sample has been obtained. Therefore, the ability to determine MGMT promoter methylation status on the basis of routine T2-w images alone is highly desirable. Additionally, because our algorithm was trained and evaluated on the multi-institutional TCIA database it is a better representative of algorithm robustness, real-world performance and potential clinical utilization than previously reported methods.^25^

The algorithm misclassified 13 cases: six subjects were misclassified as unmethylated and 7 as methylated. Despite these misclassifications, our network achieved a mean cross-validation testing accuracy of 94.73% which is higher than for the MSP, PYT, and Infinium assays.^34^ While these tissue based methods require an invasive procedure and subsequent tissue processing for at least 48 hours, our deep learning algorithm can segment the entire glioma and determine MGMT promoter methylation status in 3 minutes. The deep learning algorithm can also be fine-tuned to variations in institutional MRI scanners, while other tissue-based methods currently lacks standardization as mentioned above.

The limitations of our study are that deep learning studies require large amounts of data and the relative number of subjects with MGMT promoter methylation is small in the TCGA database. Additionally, acquisition parameters and imaging vendor platforms vary across imaging centers that contribute data. Despite these caveats our algorithm demonstrated high accuracy in determining MGMT promoter methylation status.

## CONCLUSION

We demonstrate high accuracy in determining MGMT promoter methylation status using only T2-w MR images. This represents an important milestone toward using MRI to predict glioma histology, prognosis, and appropriate treatment.

## Acknowledgments

We thank Yin Xi, PhD, statistician for help with the ROC and AUC.

