## Supplemental Material for "MRI-BASED DEEP LEARNING METHOD FOR DETERMINING METHYLATION STATUS OF THE O_6_-METHYLGUANINE–DNA METHYLTRANSFERASE PROMOTER OUTPERFORMS TISSUE BASED METHODS IN BRAIN GLIOMAS"

**FIGURES**

Supporting Figure 1: A Detailed network architecture for the MGMT-net. The previously trained 3D IDH network was used. The left arm of the Dense U-net (striped red box) is the encoder part of the network, the right arm of the network (blue box) is the decoder part and the dense block (yellow box) is the bottle neck block. The encoder part of the network was frozen to retain the pre-trained weights from the 3D IDH network. The bottleneck block and the decoder part of the network was fine-tuned for a dual class segmentation with class 1& 2 representing methylated & unmethylated MGMT promoter status respectively.

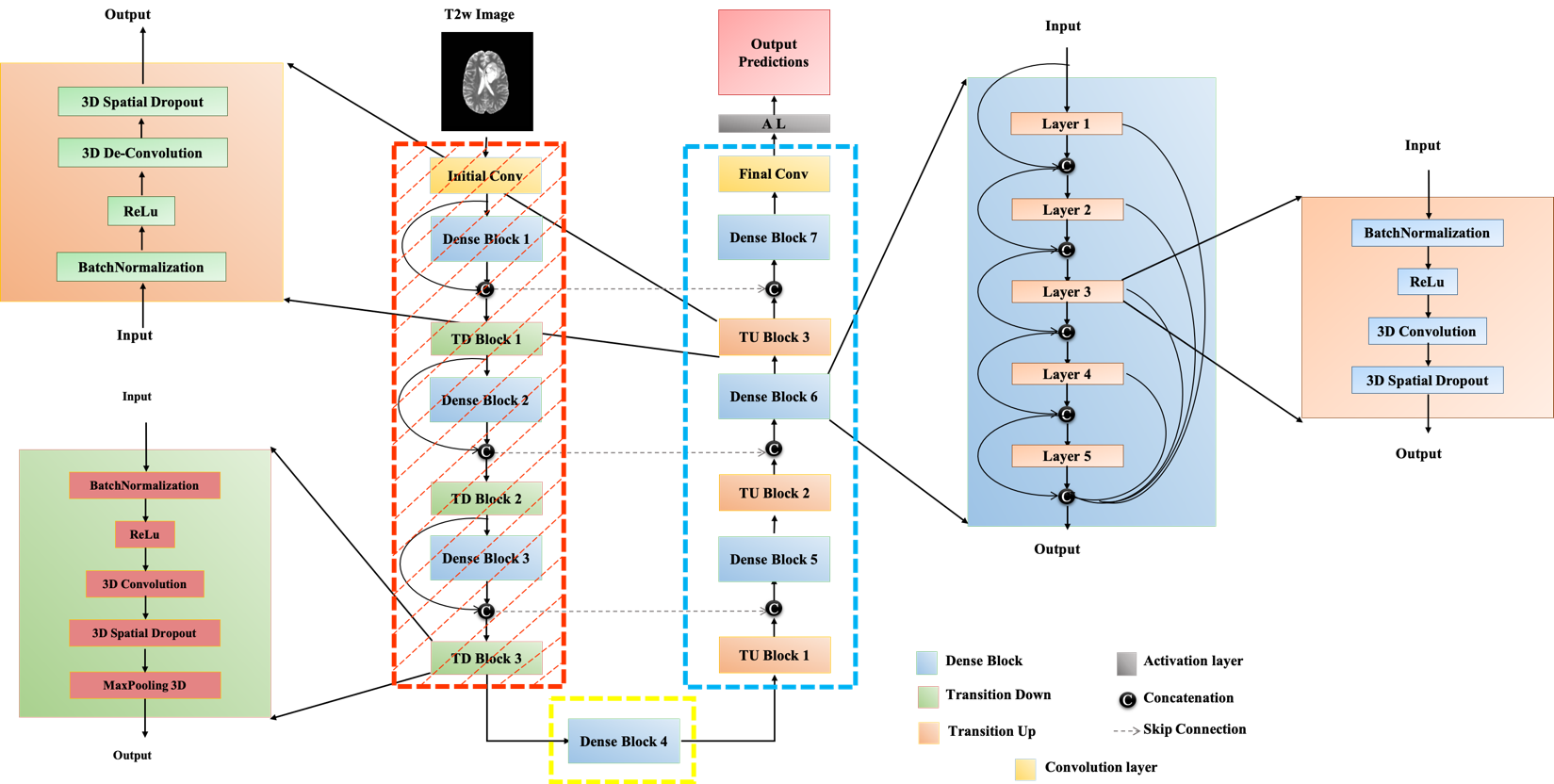

**ROC methodology**

The network output classifies voxels in the tumor as methylated or unmethylated MGMT promoter type. The percent of methylated voxels was computed for the network output for each subject in the test set by dividing the predicted methylated voxels by the total number of predicted voxels in each tumor. The percent methylated voxels can be viewed as a network output prediction likelihood of the tumor being MGMT methylated. Note that in the manuscript, majority voting (the 50% threshold) was used to determine MGMT promoter status. For the ROC analysis, the percent of methylated voxels was sorted and used as separate thresholds (cut-points) to determine the MGMT promoter status for the subjects across the test set for each new cut-point. The resulting predicted MGMT promoter class membership was compared to the ground truth values to determine sensitivity (true positive rate) and 1- specificity (false positive rate) at each threshold. The resulting values were plotted using R to obtain an ROC curve (true positive rate against false positive rate). R routines were used to fit the curves and determine the area under the curve (AUC). This procedure was repeated for each of the 3 test folds from the cross-validation procedure for the MGMT-net, providing a total of 3 ROC curves from the cross-validation.

**TABLES**

Table 1: Subject wise MGMT promoter status and tumor histology

| **Subject ID** | **Age** | **Gender** | **Histology** | **Grade** | **TCGA Data Collection** | **IDH mutation Status** | **1p/19q co-deletion status** | **MGMT promoter status** | **Cross-validation group** |
| --- | --- | --- | --- | --- | --- | --- | --- | --- | --- |
| TCGA-02-0003 | 50 | male | glioblastoma | G4 | TCGA-GBM | WT | non-codel | Unmethylated | 2 |
| TCGA-02-0006 | 56 | female | glioblastoma | G4 | TCGA-GBM | WT | non-codel | Unmethylated | 2 |
| TCGA-02-0009 | 61 | female | glioblastoma | G4 | TCGA-GBM | WT | non-codel | Unmethylated | 2 |
| TCGA-02-0011 | 18 | female | glioblastoma | G4 | TCGA-GBM | WT | non-codel | Methylated | 2 |
| TCGA-02-0027 | 33 | female | glioblastoma | G4 | TCGA-GBM | WT | non-codel | Unmethylated | 2 |
| TCGA-02-0033 | 54 | male | glioblastoma | G4 | TCGA-GBM | WT | non-codel | Methylated | 1 |
| TCGA-02-0034 | 60 | male | glioblastoma | G4 | TCGA-GBM | WT | non-codel | Unmethylated | 1 |
| TCGA-02-0037 | 74 | female | glioblastoma | G4 | TCGA-GBM | WT | non-codel | Unmethylated | 1 |
| TCGA-02-0046 | 61 | male | glioblastoma | G4 | TCGA-GBM | WT | non-codel | Methylated | 1 |
| TCGA-02-0047 | 78 | male | glioblastoma | G4 | TCGA-GBM | WT | non-codel | Unmethylated | 1 |
| TCGA-02-0060 | 66 | female | glioblastoma | G4 | TCGA-GBM | WT | non-codel | Methylated | 2 |
| TCGA-02-0064 | 50 | male | glioblastoma | G4 | TCGA-GBM | WT | non-codel | Methylated | 3 |
| TCGA-02-0069 | 31 | female | glioblastoma | G4 | TCGA-GBM | WT | non-codel | Methylated | 2 |
| TCGA-02-0075 | 63 | male | glioblastoma | G4 | TCGA-GBM | WT | non-codel | Methylated | 2 |
| TCGA-02-0086 | 45 | female | glioblastoma | G4 | TCGA-GBM | WT | non-codel | Unmethylated | 2 |
| TCGA-02-0102 | 42 | male | glioblastoma | G4 | TCGA-GBM | WT | non-codel | Unmethylated | 1 |
| TCGA-06-0119 | 81 | female | glioblastoma | G4 | TCGA-GBM | WT | non-codel | Methylated | 2 |
| TCGA-06-0122 | 84 | female | glioblastoma | G4 | TCGA-GBM | WT | non-codel | Unmethylated | 2 |
| TCGA-06-0128 | 66 | male | glioblastoma | G4 | TCGA-GBM | Mutant | non-codel | Methylated | 3 |
| TCGA-06-0129 | 30 | male | glioblastoma | G4 | TCGA-GBM | Mutant | non-codel | Methylated | 1 |
| TCGA-06-0133 | 64 | male | glioblastoma | G4 | TCGA-GBM | WT | non-codel | Unmethylated | 2 |
| TCGA-06-0137 | 63 | female | glioblastoma | G4 | TCGA-GBM | WT | non-codel | Unmethylated | 2 |
| TCGA-06-0142 | 81 | male | glioblastoma | G4 | TCGA-GBM | WT | non-codel | Unmethylated | 1 |
| TCGA-06-0143 | 58 | male | glioblastoma | G4 | TCGA-GBM | WT | non-codel | Unmethylated | 1 |
| TCGA-06-0145 | 53 | female | glioblastoma | G4 | TCGA-GBM | WT | non-codel | Methylated | 1 |
| TCGA-06-0147 | 51 | female | glioblastoma | G4 | TCGA-GBM | WT | non-codel | Methylated | 2 |
| TCGA-06-0148 | 76 | male | glioblastoma | G4 | TCGA-GBM | WT | non-codel | Unmethylated | 1 |
| TCGA-06-0881 | 50 | male | glioblastoma | G4 | TCGA-GBM | WT | non-codel | Unmethylated | 1 |
| TCGA-06-1806 | 47 | male | glioblastoma | G4 | TCGA-GBM | WT | non-codel | Unmethylated | 2 |
| TCGA-06-2570 | 21 | female | glioblastoma | G4 | TCGA-GBM | Mutant | non-codel | Methylated | 1 |
| TCGA-06-5408 | 54 | female | glioblastoma | G4 | TCGA-GBM | WT | non-codel | Unmethylated | 3 |
| TCGA-06-5412 | 78 | female | glioblastoma | G4 | TCGA-GBM | WT | non-codel | Methylated | 2 |
| TCGA-06-5413 | 67 | male | glioblastoma | G4 | TCGA-GBM | WT | non-codel | Unmethylated | 1 |
| TCGA-06-5417 | 45 | female | glioblastoma | G4 | TCGA-GBM | Mutant | NA | Methylated | 3 |
| TCGA-06-6389 | 49 | female | glioblastoma | G4 | TCGA-GBM | Mutant | non-codel | Methylated | 3 |
| TCGA-12-0829 | 75 | male | glioblastoma | G4 | TCGA-GBM | WT | non-codel | Methylated | 2 |
| TCGA-12-1093 | 66 | female | glioblastoma | G4 | TCGA-GBM | WT | non-codel | Unmethylated | 1 |
| TCGA-12-1598 | 75 | female | glioblastoma | G4 | TCGA-GBM | WT | non-codel | Methylated | 3 |
| TCGA-12-1601 | NaN | NA | NA | NA | TCGA-GBM | WT | NA | Unmethylated | 2 |
| TCGA-12-1602 | 58 | male | glioblastoma | G4 | TCGA-GBM | WT | non-codel | Methylated | 2 |
| TCGA-12-3650 | 46 | male | glioblastoma | G4 | TCGA-GBM | WT | non-codel | Unmethylated | 3 |
| TCGA-14-0789 | 54 | male | glioblastoma | G4 | TCGA-GBM | WT | non-codel | Methylated | 2 |
| TCGA-14-1456 | 23 | male | glioblastoma | G4 | TCGA-GBM | Mutant | non-codel | Unmethylated | 3 |
| TCGA-14-1794 | 59 | male | glioblastoma | G4 | TCGA-GBM | WT | non-codel | Unmethylated | 2 |
| TCGA-14-1829 | 57 | male | glioblastoma | G4 | TCGA-GBM | WT | non-codel | Unmethylated | 2 |
| TCGA-14-3477 | 38 | female | glioblastoma | G4 | TCGA-GBM | WT | non-codel | Unmethylated | 3 |
| TCGA-19-1390 | 63 | female | glioblastoma | G4 | TCGA-GBM | WT | non-codel | Methylated | 1 |
| TCGA-19-1789 | 69 | female | glioblastoma | G4 | TCGA-GBM | WT | non-codel | Methylated | 2 |
| TCGA-19-1791 | 82 | female | glioblastoma | G4 | TCGA-GBM | WT | non-codel | Unmethylated | 1 |
| TCGA-19-2620 | 70 | male | glioblastoma | G4 | TCGA-GBM | WT | non-codel | Methylated | 3 |
| TCGA-19-2624 | 51 | male | glioblastoma | G4 | TCGA-GBM | WT | non-codel | Unmethylated | 1 |
| TCGA-19-2631 | 74 | female | glioblastoma | G4 | TCGA-GBM | WT | non-codel | Methylated | 3 |
| TCGA-19-5953 | 58 | male | glioblastoma | G4 | TCGA-GBM | WT | non-codel | Methylated | 1 |
| TCGA-19-5954 | 72 | female | glioblastoma | G4 | TCGA-GBM | WT | non-codel | Methylated | 3 |
| TCGA-19-5958 | 56 | male | glioblastoma | G4 | TCGA-GBM | WT | non-codel | Unmethylated | 1 |
| TCGA-27-1830 | 57 | male | glioblastoma | G4 | TCGA-GBM | WT | non-codel | Unmethylated | 2 |
| TCGA-27-1835 | 53 | female | glioblastoma | G4 | TCGA-GBM | WT | non-codel | Methylated | 1 |
| TCGA-27-1836 | 33 | female | glioblastoma | G4 | TCGA-GBM | WT | non-codel | Methylated | 1 |
| TCGA-27-1838 | 59 | female | glioblastoma | G4 | TCGA-GBM | WT | non-codel | Unmethylated | 3 |
| TCGA-76-4925 | 76 | male | glioblastoma | G4 | TCGA-GBM | WT | non-codel | Methylated | 2 |
| TCGA-76-4926 | 68 | male | glioblastoma | G4 | TCGA-GBM | WT | non-codel | Unmethylated | 3 |
| TCGA-76-4927 | 58 | male | glioblastoma | G4 | TCGA-GBM | WT | NA | Unmethylated | 2 |
| TCGA-76-4928 | 85 | female | glioblastoma | G4 | TCGA-GBM | WT | non-codel | Methylated | 1 |
| TCGA-76-4929 | 76 | female | glioblastoma | G4 | TCGA-GBM | WT | non-codel | Methylated | 3 |
| TCGA-76-4931 | 70 | female | glioblastoma | G4 | TCGA-GBM | WT | non-codel | Unmethylated | 3 |
| TCGA-76-4932 | 50 | female | glioblastoma | G4 | TCGA-GBM | WT | NA | Methylated | 3 |
| TCGA-76-4934 | 66 | female | glioblastoma | G4 | TCGA-GBM | WT | non-codel | Methylated | 1 |
| TCGA-76-4935 | 52 | female | glioblastoma | G4 | TCGA-GBM | WT | non-codel | Methylated | 2 |
| TCGA-76-6191 | 57 | male | glioblastoma | G4 | TCGA-GBM | WT | non-codel | Unmethylated | 3 |
| TCGA-76-6192 | 74 | male | glioblastoma | G4 | TCGA-GBM | WT | non-codel | Unmethylated | 2 |
| TCGA-76-6193 | 78 | male | glioblastoma | G4 | TCGA-GBM | WT | non-codel | Unmethylated | 2 |
| TCGA-76-6280 | 57 | male | glioblastoma | G4 | TCGA-GBM | WT | non-codel | Methylated | 1 |
| TCGA-76-6282 | 63 | male | glioblastoma | G4 | TCGA-GBM | WT | non-codel | Unmethylated | 2 |
| TCGA-76-6285 | 64 | female | glioblastoma | G4 | TCGA-GBM | WT | non-codel | Unmethylated | 3 |
| TCGA-76-6286 | 60 | male | glioblastoma | G4 | TCGA-GBM | WT | non-codel | Unmethylated | 1 |
| TCGA-76-6656 | 66 | male | glioblastoma | G4 | TCGA-GBM | WT | non-codel | Methylated | 3 |
| TCGA-76-6657 | 74 | male | glioblastoma | G4 | TCGA-GBM | WT | non-codel | Methylated | 2 |
| TCGA-76-6661 | 54 | male | glioblastoma | G4 | TCGA-GBM | WT | non-codel | Unmethylated | 3 |
| TCGA-76-6662 | 58 | male | glioblastoma | G4 | TCGA-GBM | WT | non-codel | Unmethylated | 1 |
| TCGA-76-6663 | 44 | female | glioblastoma | G4 | TCGA-GBM | WT | non-codel | Unmethylated | 3 |
| TCGA-76-6664 | 49 | female | glioblastoma | G4 | TCGA-GBM | WT | non-codel | Methylated | 3 |
| TCGA-CS-4938 | 31 | female | astrocytoma | G2 | TCGA-LGG | Mutant | non-codel | Unmethylated | 3 |
| TCGA-CS-4941 | 67 | male | astrocytoma | G3 | TCGA-LGG | WT | non-codel | Methylated | 1 |
| TCGA-CS-4942 | 44 | female | astrocytoma | G3 | TCGA-LGG | Mutant | non-codel | Unmethylated | 1 |
| TCGA-CS-4943 | 37 | male | astrocytoma | G3 | TCGA-LGG | Mutant | non-codel | Methylated | 3 |
| TCGA-CS-4944 | 50 | male | astrocytoma | G2 | TCGA-LGG | Mutant | non-codel | Methylated | 2 |
| TCGA-CS-5390 | 47 | female | oligodendroglioma | G2 | TCGA-LGG | Mutant | codel | Methylated | 2 |
| TCGA-CS-5393 | 39 | male | astrocytoma | G3 | TCGA-LGG | Mutant | non-codel | Methylated | 2 |
| TCGA-CS-5394 | 40 | male | astrocytoma | G3 | TCGA-LGG | Mutant | non-codel | Methylated | 1 |
| TCGA-CS-5395 | 43 | male | oligodendroglioma | G2 | TCGA-LGG | WT | non-codel | Unmethylated | 1 |
| TCGA-CS-5396 | 53 | female | oligodendroglioma | G3 | TCGA-LGG | Mutant | codel | Methylated | 3 |
| TCGA-CS-5397 | 54 | female | astrocytoma | G3 | TCGA-LGG | WT | non-codel | Unmethylated | 2 |
| TCGA-CS-6186 | 58 | male | oligoastrocytoma | G3 | TCGA-LGG | WT | non-codel | Unmethylated | 1 |
| TCGA-CS-6188 | 48 | male | astrocytoma | G3 | TCGA-LGG | WT | non-codel | Unmethylated | 2 |
| TCGA-CS-6290 | 31 | male | astrocytoma | G3 | TCGA-LGG | Mutant | non-codel | Methylated | 1 |
| TCGA-CS-6665 | 51 | female | astrocytoma | G3 | TCGA-LGG | Mutant | non-codel | Methylated | 3 |
| TCGA-CS-6666 | 22 | male | astrocytoma | G3 | TCGA-LGG | Mutant | non-codel | Methylated | 2 |
| TCGA-CS-6667 | 39 | female | astrocytoma | G2 | TCGA-LGG | Mutant | non-codel | Methylated | 1 |
| TCGA-CS-6668 | 57 | female | oligodendroglioma | G2 | TCGA-LGG | Mutant | codel | Methylated | 1 |
| TCGA-CS-6669 | 26 | female | oligodendroglioma | G2 | TCGA-LGG | WT | non-codel | Unmethylated | 3 |
| TCGA-DU-5849 | 48 | male | oligodendroglioma | G2 | TCGA-LGG | Mutant | codel | Methylated | 1 |
| TCGA-DU-5851 | 40 | female | oligoastrocytoma | G3 | TCGA-LGG | Mutant | non-codel | Unmethylated | 3 |
| TCGA-DU-5852 | 61 | female | oligoastrocytoma | G3 | TCGA-LGG | WT | non-codel | Methylated | 3 |
| TCGA-DU-5853 | 29 | male | oligoastrocytoma | G2 | TCGA-LGG | Mutant | non-codel | Methylated | 2 |
| TCGA-DU-5854 | 57 | female | astrocytoma | G3 | TCGA-LGG | WT | non-codel | Unmethylated | 2 |
| TCGA-DU-5855 | 49 | female | oligoastrocytoma | G3 | TCGA-LGG | Mutant | non-codel | Methylated | 2 |
| TCGA-DU-5871 | 37 | female | oligoastrocytoma | G2 | TCGA-LGG | Mutant | non-codel | Methylated | 3 |
| TCGA-DU-5872 | 43 | female | oligoastrocytoma | G2 | TCGA-LGG | Mutant | non-codel | Methylated | 2 |
| TCGA-DU-5874 | 62 | female | oligodendroglioma | G2 | TCGA-LGG | Mutant | codel | Methylated | 3 |
| TCGA-DU-6395 | 31 | male | oligoastrocytoma | G2 | TCGA-LGG | Mutant | non-codel | Methylated | 1 |
| TCGA-DU-6397 | 45 | male | oligodendroglioma | G3 | TCGA-LGG | Mutant | codel | Methylated | 2 |
| TCGA-DU-6399 | 54 | male | oligodendroglioma | G2 | TCGA-LGG | Mutant | non-codel | Methylated | 1 |
| TCGA-DU-6400 | 66 | female | oligodendroglioma | G2 | TCGA-LGG | Mutant | codel | Methylated | 3 |
| TCGA-DU-6401 | 31 | female | oligodendroglioma | G2 | TCGA-LGG | Mutant | non-codel | Methylated | 3 |
| TCGA-DU-6404 | 24 | female | oligodendroglioma | G3 | TCGA-LGG | WT | non-codel | Unmethylated | 3 |
| TCGA-DU-6405 | 51 | female | astrocytoma | G3 | TCGA-LGG | WT | non-codel | Methylated | 3 |
| TCGA-DU-6407 | 35 | female | oligodendroglioma | G2 | TCGA-LGG | Mutant | non-codel | Methylated | 3 |
| TCGA-DU-6408 | 23 | female | oligodendroglioma | G3 | TCGA-LGG | Mutant | non-codel | Methylated | 1 |
| TCGA-DU-7008 | 41 | female | oligodendroglioma | G2 | TCGA-LGG | Mutant | non-codel | Methylated | 1 |
| TCGA-DU-7010 | 58 | female | astrocytoma | G3 | TCGA-LGG | Mutant | non-codel | Methylated | 3 |
| TCGA-DU-7013 | 59 | male | astrocytoma | G3 | TCGA-LGG | WT | non-codel | Unmethylated | 1 |
| TCGA-DU-7015 | 41 | female | oligodendroglioma | G2 | TCGA-LGG | Mutant | non-codel | Methylated | 2 |
| TCGA-DU-7018 | 57 | female | oligodendroglioma | G3 | TCGA-LGG | Mutant | codel | Methylated | 1 |
| TCGA-DU-7019 | 39 | male | oligoastrocytoma | G3 | TCGA-LGG | Mutant | non-codel | Methylated | 1 |
| TCGA-DU-7294 | 53 | female | oligodendroglioma | G2 | TCGA-LGG | Mutant | codel | Methylated | 1 |
| TCGA-DU-7298 | 38 | female | astrocytoma | G3 | TCGA-LGG | Mutant | non-codel | Methylated | 2 |
| TCGA-DU-7299 | 33 | male | astrocytoma | G3 | TCGA-LGG | Mutant | non-codel | Methylated | 3 |
| TCGA-DU-7300 | 53 | female | oligodendroglioma | G3 | TCGA-LGG | Mutant | codel | Methylated | 3 |
| TCGA-DU-7301 | 53 | male | oligodendroglioma | G2 | TCGA-LGG | Mutant | non-codel | Methylated | 3 |
| TCGA-DU-7302 | 48 | female | oligodendroglioma | G3 | TCGA-LGG | Mutant | codel | Methylated | 1 |
| TCGA-DU-7304 | 43 | male | oligoastrocytoma | G3 | TCGA-LGG | Mutant | non-codel | Methylated | 3 |
| TCGA-DU-7306 | 67 | male | oligoastrocytoma | G2 | TCGA-LGG | Mutant | non-codel | Methylated | 1 |
| TCGA-DU-7309 | 41 | female | oligodendroglioma | G3 | TCGA-LGG | Mutant | non-codel | Methylated | 3 |
| TCGA-DU-8158 | 57 | female | astrocytoma | G3 | TCGA-LGG | WT | non-codel | Unmethylated | 3 |
| TCGA-DU-8162 | 61 | female | oligoastrocytoma | G3 | TCGA-LGG | WT | non-codel | Unmethylated | 1 |
| TCGA-DU-8164 | 51 | male | oligodendroglioma | G2 | TCGA-LGG | Mutant | codel | Methylated | 1 |
| TCGA-DU-8165 | 60 | female | oligodendroglioma | G3 | TCGA-LGG | WT | non-codel | Unmethylated | 1 |
| TCGA-DU-8166 | 29 | female | oligoastrocytoma | G2 | TCGA-LGG | Mutant | non-codel | Methylated | 2 |
| TCGA-DU-8167 | 69 | female | oligoastrocytoma | G2 | TCGA-LGG | Mutant | non-codel | Methylated | 3 |
| TCGA-DU-8168 | 55 | female | oligodendroglioma | G3 | TCGA-LGG | Mutant | codel | Methylated | 3 |
| TCGA-DU-A5TP | 33 | male | astrocytoma | G3 | TCGA-LGG | Mutant | non-codel | Methylated | 2 |
| TCGA-DU-A5TR | 51 | male | oligoastrocytoma | G2 | TCGA-LGG | Mutant | non-codel | Methylated | 2 |
| TCGA-DU-A5TS | 42 | male | oligodendroglioma | G2 | TCGA-LGG | Mutant | non-codel | Methylated | 2 |
| TCGA-DU-A5TT | 70 | male | oligodendroglioma | G3 | TCGA-LGG | WT | non-codel | Methylated | 2 |
| TCGA-DU-A5TU | 62 | female | astrocytoma | G2 | TCGA-LGG | Mutant | non-codel | Methylated | 1 |
| TCGA-DU-A5TW | 33 | female | astrocytoma | G3 | TCGA-LGG | Mutant | non-codel | Methylated | 2 |
| TCGA-DU-A5TY | 46 | female | astrocytoma | G3 | TCGA-LGG | WT | non-codel | Methylated | 2 |
| TCGA-DU-A6S2 | 37 | female | oligodendroglioma | G2 | TCGA-LGG | Mutant | codel | Methylated | 2 |
| TCGA-DU-A6S3 | 60 | male | oligodendroglioma | G2 | TCGA-LGG | Mutant | codel | Methylated | 3 |
| TCGA-DU-A6S6 | 35 | female | oligoastrocytoma | G2 | TCGA-LGG | Mutant | codel | Methylated | 1 |
| TCGA-DU-A6S7 | 27 | female | astrocytoma | G3 | TCGA-LGG | Mutant | non-codel | Methylated | 2 |
| TCGA-DU-A6S8 | 74 | female | oligodendroglioma | G3 | TCGA-LGG | Mutant | codel | Methylated | 2 |
| TCGA-FG-5963 | 23 | male | astrocytoma | G3 | TCGA-LGG | WT | non-codel | Unmethylated | 2 |
| TCGA-FG-5964 | 62 | male | oligodendroglioma | G2 | TCGA-LGG | Mutant | codel | Methylated | 1 |
| TCGA-FG-6688 | 59 | female | astrocytoma | G3 | TCGA-LGG | WT | non-codel | Methylated | 2 |
| TCGA-FG-6689 | 30 | male | astrocytoma | G2 | TCGA-LGG | Mutant | non-codel | Methylated | 3 |
| TCGA-FG-6690 | 70 | male | oligodendroglioma | G2 | TCGA-LGG | Mutant | non-codel | Methylated | 2 |
| TCGA-FG-6691 | 23 | female | astrocytoma | G2 | TCGA-LGG | Mutant | non-codel | Unmethylated | 3 |
| TCGA-FG-6692 | 63 | male | oligodendroglioma | G3 | TCGA-LGG | WT | non-codel | Methylated | 1 |
| TCGA-FG-7634 | 28 | male | oligodendroglioma | G2 | TCGA-LGG | Mutant | codel | Methylated | 1 |
| TCGA-FG-7637 | 49 | male | oligoastrocytoma | G2 | TCGA-LGG | Mutant | non-codel | Methylated | 1 |
| TCGA-FG-8189 | 33 | female | oligodendroglioma | G2 | TCGA-LGG | Mutant | non-codel | Methylated | 3 |
| TCGA-FG-A4MT | 27 | female | oligodendroglioma | G2 | TCGA-LGG | Mutant | non-codel | Methylated | 2 |
| TCGA-FG-A4MU | 58 | male | oligoastrocytoma | G3 | TCGA-LGG | WT | non-codel | Methylated | 1 |
| TCGA-FG-A6IZ | 60 | male | oligodendroglioma | G2 | TCGA-LGG | Mutant | codel | Methylated | 3 |
| TCGA-FG-A6J1 | 44 | female | oligodendroglioma | G2 | TCGA-LGG | Mutant | codel | Methylated | 2 |
| TCGA-FG-A713 | 74 | female | oligoastrocytoma | G2 | TCGA-LGG | Mutant | codel | Methylated | 3 |
| TCGA-FG-A87N | 37 | female | astrocytoma | G3 | TCGA-LGG | Mutant | non-codel | Methylated | 3 |
| TCGA-HT-7468 | 30 | male | oligodendroglioma | G3 | TCGA-LGG | Mutant | codel | Methylated | 1 |
| TCGA-HT-7469 | 30 | male | oligodendroglioma | G3 | TCGA-LGG | WT | non-codel | Methylated | 1 |
| TCGA-HT-7471 | 37 | female | oligodendroglioma | G3 | TCGA-LGG | Mutant | codel | Methylated | 2 |
| TCGA-HT-7472 | 38 | male | oligodendroglioma | G2 | TCGA-LGG | Mutant | non-codel | Methylated | 1 |
| TCGA-HT-7473 | 28 | male | oligoastrocytoma | G2 | TCGA-LGG | Mutant | non-codel | Unmethylated | 3 |
| TCGA-HT-7475 | 67 | male | oligoastrocytoma | G3 | TCGA-LGG | Mutant | non-codel | Methylated | 3 |
| TCGA-HT-7476 | 26 | male | astrocytoma | G2 | TCGA-LGG | Mutant | non-codel | Methylated | 1 |
| TCGA-HT-7478 | 36 | male | astrocytoma | G2 | TCGA-LGG | Mutant | non-codel | Unmethylated | 1 |
| TCGA-HT-7481 | 39 | male | oligodendroglioma | G2 | TCGA-LGG | Mutant | codel | Methylated | 2 |
| TCGA-HT-7602 | 21 | male | oligodendroglioma | G2 | TCGA-LGG | Mutant | non-codel | Methylated | 2 |
| TCGA-HT-7603 | 29 | male | oligodendroglioma | G2 | TCGA-LGG | Mutant | non-codel | Methylated | 2 |
| TCGA-HT-7605 | 38 | male | oligodendroglioma | G2 | TCGA-LGG | Mutant | codel | Methylated | 3 |
| TCGA-HT-7606 | 30 | female | astrocytoma | G2 | TCGA-LGG | Mutant | non-codel | Unmethylated | 1 |
| TCGA-HT-7608 | 61 | male | oligoastrocytoma | G2 | TCGA-LGG | Mutant | codel | Methylated | 2 |
| TCGA-HT-7616 | 75 | male | oligodendroglioma | G3 | TCGA-LGG | Mutant | codel | Methylated | 3 |
| TCGA-HT-7680 | 32 | female | astrocytoma | G2 | TCGA-LGG | WT | non-codel | Unmethylated | 3 |
| TCGA-HT-7684 | 58 | male | oligoastrocytoma | G3 | TCGA-LGG | Mutant | non-codel | Methylated | 3 |
| TCGA-HT-7686 | 29 | female | astrocytoma | G3 | TCGA-LGG | Mutant | non-codel | Methylated | 3 |
| TCGA-HT-7690 | 29 | male | oligoastrocytoma | G3 | TCGA-LGG | Mutant | non-codel | Methylated | 3 |
| TCGA-HT-7692 | 43 | male | oligoastrocytoma | G2 | TCGA-LGG | Mutant | codel | Methylated | 1 |
| TCGA-HT-7693 | 51 | female | oligodendroglioma | G2 | TCGA-LGG | Mutant | non-codel | Methylated | 1 |
| TCGA-HT-7694 | 60 | male | oligodendroglioma | G3 | TCGA-LGG | Mutant | codel | Methylated | 2 |
| TCGA-HT-7695 | 29 | female | oligodendroglioma | G2 | TCGA-LGG | Mutant | codel | Methylated | 1 |
| TCGA-HT-7854 | 62 | male | astrocytoma | G2 | TCGA-LGG | WT | non-codel | Unmethylated | 3 |
| TCGA-HT-7855 | 39 | male | astrocytoma | G3 | TCGA-LGG | Mutant | non-codel | Methylated | 2 |
| TCGA-HT-7856 | 35 | male | oligodendroglioma | G3 | TCGA-LGG | Mutant | codel | Methylated | 2 |
| TCGA-HT-7860 | 60 | female | astrocytoma | G3 | TCGA-LGG | WT | non-codel | Methylated | 2 |
| TCGA-HT-7874 | 41 | female | oligodendroglioma | G3 | TCGA-LGG | Mutant | codel | Methylated | 1 |
| TCGA-HT-7877 | 20 | female | oligodendroglioma | G2 | TCGA-LGG | Mutant | codel | Methylated | 3 |
| TCGA-HT-7879 | 31 | male | oligoastrocytoma | G3 | TCGA-LGG | Mutant | non-codel | Methylated | 1 |
| TCGA-HT-7880 | 30 | male | oligoastrocytoma | G2 | TCGA-LGG | Mutant | non-codel | Methylated | 3 |
| TCGA-HT-7882 | 66 | male | oligodendroglioma | G3 | TCGA-LGG | WT | non-codel | Methylated | 2 |
| TCGA-HT-7884 | 44 | female | astrocytoma | G2 | TCGA-LGG | Mutant | non-codel | Methylated | 1 |
| TCGA-HT-7902 | 30 | female | oligoastrocytoma | G2 | TCGA-LGG | Mutant | non-codel | Methylated | 3 |
| TCGA-HT-8010 | 64 | female | oligodendroglioma | G2 | TCGA-LGG | Mutant | codel | Methylated | 3 |
| TCGA-HT-8013 | 37 | female | oligoastrocytoma | G2 | TCGA-LGG | Mutant | non-codel | Methylated | 1 |
| TCGA-HT-8015 | 21 | male | astrocytoma | G2 | TCGA-LGG | WT | non-codel | Unmethylated | 1 |
| TCGA-HT-8018 | 40 | female | oligoastrocytoma | G2 | TCGA-LGG | Mutant | non-codel | Methylated | 3 |
| TCGA-HT-8019 | 34 | female | oligodendroglioma | G3 | TCGA-LGG | WT | non-codel | Unmethylated | 1 |
| TCGA-HT-8105 | 54 | male | oligodendroglioma | G3 | TCGA-LGG | Mutant | codel | Methylated | 3 |
| TCGA-HT-8106 | 53 | male | astrocytoma | G3 | TCGA-LGG | Mutant | non-codel | Methylated | 1 |
| TCGA-HT-8107 | 62 | male | oligodendroglioma | G2 | TCGA-LGG | WT | non-codel | Methylated | 2 |
| TCGA-HT-8108 | 26 | female | oligodendroglioma | G2 | TCGA-LGG | Mutant | non-codel | Methylated | 1 |
| TCGA-HT-8111 | 32 | male | oligoastrocytoma | G3 | TCGA-LGG | Mutant | non-codel | Methylated | 2 |
| TCGA-HT-8113 | 49 | female | oligodendroglioma | G2 | TCGA-LGG | Mutant | non-codel | Methylated | 3 |
| TCGA-HT-8114 | 36 | male | oligoastrocytoma | G3 | TCGA-LGG | Mutant | non-codel | Methylated | 2 |
| TCGA-HT-8558 | 29 | female | oligodendroglioma | G2 | TCGA-LGG | WT | non-codel | Unmethylated | 2 |
| TCGA-HT-8563 | 30 | female | astrocytoma | G3 | TCGA-LGG | Mutant | non-codel | Unmethylated | 3 |
| TCGA-HT-8564 | 47 | male | astrocytoma | G3 | TCGA-LGG | WT | non-codel | Unmethylated | 2 |
| TCGA-HT-A4DS | 55 | female | astrocytoma | G3 | TCGA-LGG | WT | non-codel | Unmethylated | 2 |
| TCGA-HT-A5R5 | 33 | female | oligodendroglioma | G2 | TCGA-LGG | Mutant | non-codel | Methylated | 2 |
| TCGA-HT-A5RB | 24 | male | astrocytoma | G2 | TCGA-LGG | Mutant | non-codel | Methylated | 1 |
| TCGA-HT-A5RC | 70 | female | astrocytoma | G3 | TCGA-LGG | WT | non-codel | Unmethylated | 2 |
| TCGA-HT-A616 | 36 | female | astrocytoma | G2 | TCGA-LGG | Mutant | non-codel | Methylated | 3 |
| TCGA-HT-A61A | 20 | female | oligodendroglioma | G2 | TCGA-LGG | Mutant | non-codel | Methylated | 3 |
| TCGA-HT-A61B | NaN | NaN | NaN | NaN | TCGA-LGG | Mutant | non-codel | Methylated | 2 |
| W1_19961025 | NaN | NaN | NaN | NaN | NaN | NaN | NaN | Unmethylated | 2 |
| W10_19970429 | NaN | NaN | NaN | NaN | NaN | NaN | NaN | Methylated | 1 |
| W12_19970620 | NaN | NaN | NaN | NaN | NaN | NaN | NaN | Unmethylated | 3 |
| W13_19970822 | NaN | NaN | NaN | NaN | NaN | NaN | NaN | Unmethylated | 3 |
| W16_19971015 | NaN | NaN | NaN | NaN | NaN | NaN | NaN | Unmethylated | 1 |
| W18_19971110 | NaN | NaN | NaN | NaN | NaN | NaN | NaN | Methylated | 1 |
| W2_19961101 | NaN | NaN | NaN | NaN | NaN | NaN | NaN | Methylated | 3 |
| W20_19970516 | NaN | NaN | NaN | NaN | NaN | NaN | NaN | Unmethylated | 1 |
| W21_19980105 | NaN | NaN | NaN | NaN | NaN | NaN | NaN | Unmethylated | 2 |
| W22_19980102 | NaN | NaN | NaN | NaN | NaN | NaN | NaN | Methylated | 1 |
| W29_19980521 | NaN | NaN | NaN | NaN | NaN | NaN | NaN | Unmethylated | 3 |
| W30_19980608 | NaN | NaN | NaN | NaN | NaN | NaN | NaN | Methylated | 2 |
| W31_19980629 | NaN | NaN | NaN | NaN | NaN | NaN | NaN | Unmethylated | 3 |
| W32_19980701 | NaN | NaN | NaN | NaN | NaN | NaN | NaN | Methylated | 3 |
| W33_19980704 | NaN | NaN | NaN | NaN | NaN | NaN | NaN | Methylated | 3 |
| W34_19980713 | NaN | NaN | NaN | NaN | NaN | NaN | NaN | Unmethylated | 2 |
| W36_19980714 | NaN | NaN | NaN | NaN | NaN | NaN | NaN | Unmethylated | 1 |
| W38_19980910 | NaN | NaN | NaN | NaN | NaN | NaN | NaN | Methylated | 2 |
| W39_19980919 | NaN | NaN | NaN | NaN | NaN | NaN | NaN | Methylated | 1 |
| W5_19961211 | NaN | NaN | NaN | NaN | NaN | NaN | NaN | Unmethylated | 3 |
| W54_20000902 | NaN | NaN | NaN | NaN | NaN | NaN | NaN | Unmethylated | 3 |
| W7_19961218 | NaN | NaN | NaN | NaN | NaN | NaN | NaN | Methylated | 1 |
| W9_19970410 | NaN | NaN | NaN | NaN | NaN | NaN | NaN | Unmethylated | 3 |
